# microRNA-dependent control of sensory neuron function regulates posture behaviour in *Drosophila*

**DOI:** 10.1101/2020.08.24.262626

**Authors:** M. Klann, A.R. Issa, C.R. Alonso

## Abstract

All what we see, touch, hear, taste or smell must first be detected by the sensory elements in our nervous system. Sensory neurons, therefore, represent a critical component in all neural circuits and their correct function is essential for the generation of behaviour and adaptation to the environment. Here we report that a gene encoding the evolutionarily conserved microRNA (miRNA) *miR-263b*, plays a key behavioural role in *Drosophila* through effects on the function of larval sensory neurons. Several independent experiments support this finding. First, miRNA expression analysis by means of a *miR-263b* reporter line, and fluorescent-activated cell sorting coupled to quantitative PCR, both demonstrate expression of *miR-263b* in *Drosophila* larval sensory neurons. Second, behavioural tests in *miR-263b* null mutants show defects in self-righting: an innate and evolutionarily conserved posture control behaviour that allows the larva to return to its normal position if turned upside-down. Third, competitive inhibition of *miR-263b* in sensory neurons using a *miR-263b* ‘sponge’ leads to self-righting defects. Fourth, systematic analysis of sensory neurons in *miR-263b* mutants shows no detectable morphological defects in their stereotypic pattern. Fifth, genetically-encoded calcium sensors expressed in the sensory domain reveal a reduction in neural activity in *miR-263b* null mutants. Sixth, *miR-263b* null mutants show a reduced ‘touch-response’ behaviour and a compromised response to sound, both characteristic of larval sensory deficits. Furthermore, bioinformatic miRNA target analysis, gene expression assays, and behavioural phenocopy experiments suggest that *miR-263b* might exert its effects – at least in part – through repression of the bHLH transcription factor *atonal*. Altogether, our study suggests a model in which miRNA-dependent control of transcription factor expression affects sensory function and behaviour. Building on the evolutionary conservation of *miR-263b*, we propose that similar processes may modulate sensory function in other animals, including mammals.

## INTRODUCTION

Although mechanisms of biological communication – i.e. effective delivery of information – are fundamental to biological processes across all scales and species (Lerner et al., 2016), the nervous system is probably one of the best examples of high-speed and complex information transmitted within a cellular network.

In neural systems, input information is represented by sensory signals which provide the brain with essential information about the external environment, so that adequate actions can be selected and implemented. Sensory information is encoded in the form of firing patterns of populations of peripheral neurons, collectively known as sensory neurons. To recognise minor – yet, potentially crucial – changes in the external world, sensory neurons must detect a range of diverse stimuli, and transform them into sequences of neuronal activity, which can subsequently be transmitted to the rest of the system. This requires that neural network components are ‘tuned’ or aligned by biochemical machineries operating through a common language, so that neurons can talk to one another in an efficient manner, at rates compatible with the speed of the behaviours they control. In fact, the physiology of neurons stems from coherent gene regulatory systems able to balance neuronal stoichiometry: ion-channels are composed of several protein subunits, neurotransmitter synthesis requires enzymatic pathways with multiple components, and transmitter release entails complex machineries made of proteins from different families (Davies, 2004); all these elements require dynamic qualitative and quantitative feed-back control systems able to monitor protein concentrations and be capable to maintain expression levels within a suitable range. These molecular control devices rely on both, transcriptional as well as post-transcriptional processes.

In this paper we focus on the post-transcriptional component, studying the roles played by small regulatory non-coding RNAs termed microRNAs (miRNAs) on the specification of sensory neurons in *Drosophila* larvae. miRNAs regulate the expression of a suite of protein encoding mRNAs by inducing their degradation and/or blocking their translation into protein (Bartel, 2018). Removal of miRNA genes can lead to target de-repression (up-regulation), and this, may, in certain circumstances, disrupt neural functions critical to physiology and behavioural control.

Our focus on miRNA roles is based on a recent discovery (in our group) that mutation of a single miRNA gene in *Drosophila* disrupts a particular larval locomotor behaviour termed *self-righting*: a movement that restores normal position after the animal is placed upside-down (Picao-Osorio et al., 2015). Mapping the ‘focus’ of action (Benzer, 1967; Hotta and Benzer, 1972) of this miRNA (*miR-iab4*) within the fruit fly larva led to the finding that this gene did not affect neural development; instead, it modifies the physiology of a set of motor neurons, innervating lateral transverse muscles 1 and 2 (LT1/2-MNs) in the *Drosophila* larva (Picao-Osorio et al., 2015). Extensions of this work to later stages of fly development, demonstrated that *miR-iab4* also controls self-righting in the *Drosophila* adult (Issa et al., 2019), through actions on a different set of motor neurons indicating that individual miRNA molecules may affect equivalent behaviours on systems bearing profoundly different anatomy, neurophysiology and biomechanics. Furthermore, a genetic screen in fruit fly larvae (Picao-Osorio et al., 2017) showed that multiple miRNAs can affect self-righting, revealing a pervasive influence of miRNA control on this postural behaviour.

Self-righting is a complex evolutionarily conserved three-dimensional adaptive and innate locomotor sequence (Ashe, 1970; Faisal and Matheson, 2001; Jusufi et al., 2011). To trigger this movement, the fruit fly larva must, first, determine that its position is abnormal, suggesting that sensory processes might play a key role in this adaptive behaviour. *Drosophila* larvae possess a wide range of sensory organs including multidendritic (md) SNs (Bodmer and Jan, 1987; Grueber et al., 2002) and chordotonal organs (Field and Matheson, 1998) capable of detecting chemical and mechanical inputs. These sensory systems are present on the body wall, arranged in highly stereotypical patterns within each segment of the larva, and show complex morphologies (Zawarzin, 1912; Hartenstein, 1988).

Here we show that the evolutionarily conserved miRNA *miR-263b* is essential for larval self-righting, through modulatory effects on the sensory system. Cell ablation experiments show that sensory neurons are essential for self-righting, and gene expression analyses demonstrate that *miR-263b* is expressed across different populations of larval sensory neurons. Genetic manipulations show that normal *miR-263b* expression in these sensory elements is essential for normal self-righting; furthermore, detailed morphological analyses of the sensory system demonstrate that lack of *miR-263b* does not disrupt the intricate morphology or array of larval sensory neurons, suggesting that this miRNA might, instead, affect the function of these neurons. Behavioural and optical imaging experiments confirm this, indicating that the functionality and physiology of sensory neurons is abnormal in the absence of *miR-263b*. Lastly, based on bioinformatic, gene expression and behavioural analyses, we propose a model in which *miR-263b* exerts its actions on sensory neurons via repression of the bHLH transcription factor *atonal*. Our work thus provides evidence that miRNAs can control complex motor behaviours by modulating the physiology of sensory neurons.

## RESULTS

### Sensory and genetic requirements for self-righting behaviour

The self-righting response is an innate and evolutionarily conserved movement that involves body rotation when the organism is placed upside-down (Ashe, 1970; Faisal and Matheson, 2001; Jusufi et al., 2011; Picao-Osorio et al., 2015). In *Drosophila* larvae, self-righting concerns the coordinated tri-dimensional motion of multiple larval segments (**Figure 1A-B**) and is triggered by the inversion of the position of the larval body in respect to the substrate. A possibility to explain the activation of self-righting is that the response might be triggered by a change in the orientation of the gravitational field; but this is not the case: inversion of the polarity of the gravitational field does not affect larval movement or trigger the self-righting response (Movies S1-S2). An alternative model is that the animal detects an anomaly in the normal pattern of sensory stimuli that informs the status of its contact with substrate and, based on this change, triggers the self-righting sequence. Peripheral sensory inputs are detected by genetically-defined subsets of larval sensory neurons arranged dorsoventrally along the larval body wall (Zawarzin A., 1912; Hartenstein, 1988), including: multidendritic (md) sensory neurons (Bodmer and Jan, 1987; Grueber et al., 2002) and chordotonal organs (Field and Matheson, 1998) (**Figure 1C-E**). We hypothesised that these peripheral sensors may play a role in conveying the necessary information that triggers self-righting and, to test this model, we disabled subsets of peripheral SNs using the tetanus toxin (Sweeney et al., 1995) and examined the effects of these perturbations on self-righting behaviour. The results of these experiments demonstrate that SNs demarked by expression of the drivers *109(2)80* (multiple dendritic neurons, oenocytes and chordotonal organs (Gao et al., 1999)), *ppk* (class IV dendritic arborization neurons and, less strongly, class III neurons (Grueber et al., 2002; Grueber et al., 2007) and *iav* (chordotonal neurons (Kwon et al., 2010) are essential for normal self-righting (**Figure 1F**).

**Figure 1.**
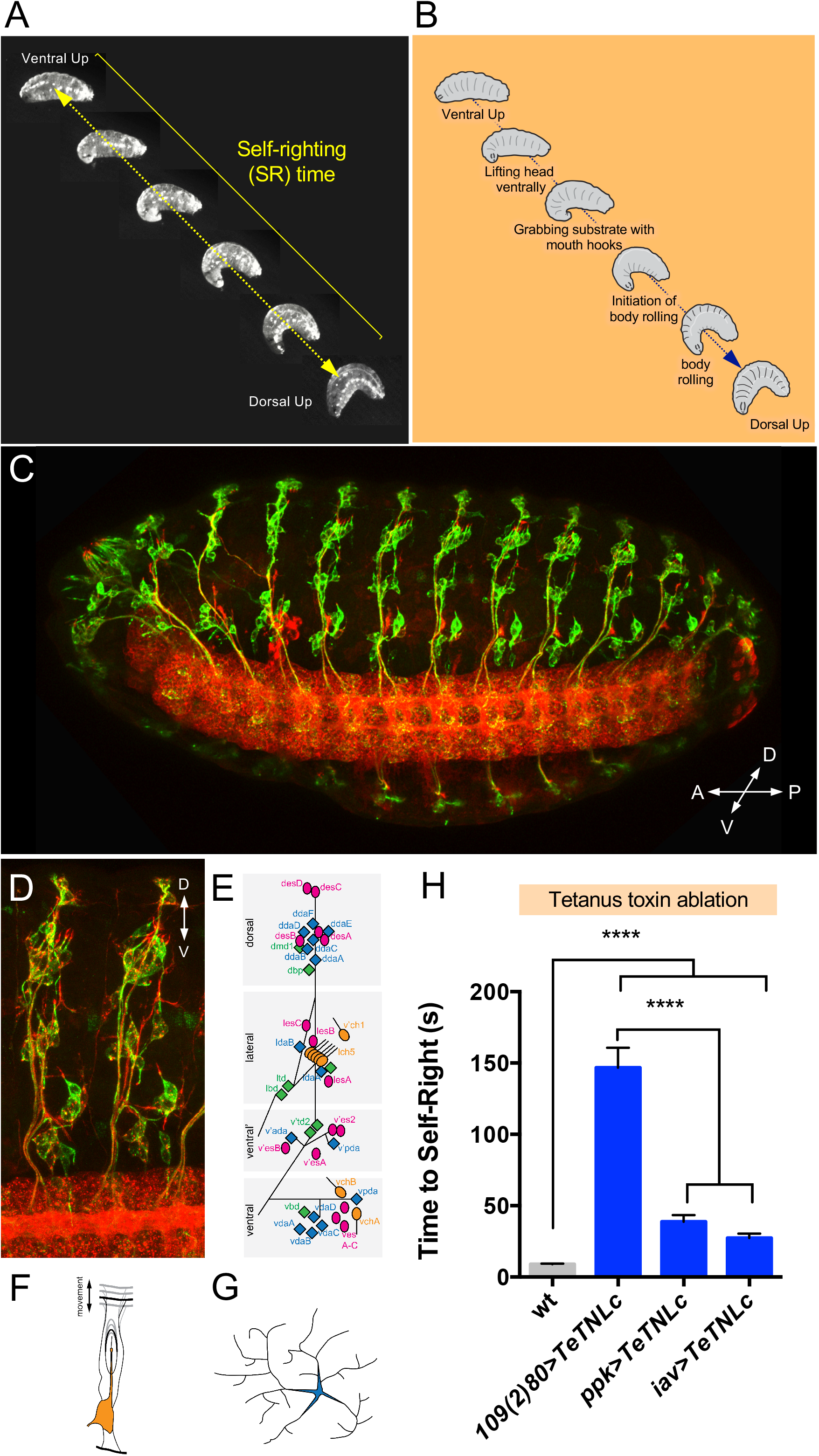
Self-righting behaviour and its sensory requirements. **(A)**. The self-righting sequence shown as a series of still images acquired from a video captured during larval self-righting (arrow, self-righting time) **(B)** Self-righting sequence. Schematic representation indicating the individual steps observed during the self-righting behavioural sequence. **(C)** The sensory system of stage 16 Drosophila embryo, in ventro-lateral view. Sensory neurons were labelled with α-22C10 (green) (α-HRP (red) provides a general neuronal marker). **(D)**. Sensory organ arrangement of embryonic abdominal segments, ventro-lateral view. Note the complexity and regularity of the arrangement of elements. **(E)**. Schematic representation of the distribution and composition of embryonic/larval sensory organs in the abdomen; **(F)** Morphology of chordotonal organs, and **(G)** multidendritic neurons is depicted by means of cartoons. **(H)** Self-righting performance is decreased when all components of the sensory system (109(2)80) or some of them (ppk, multidendritic SNs; iav, chordotonal organs) are disabled by the tetanus toxin, indicating that information detected by the sensory system is essential for normal larval self-righting.

### Genetic elements affecting self-righting sensory function: a role for small non-coding RNAs

The physiology of SNs (and that of all neurons) largely relies on biochemistry; and the latter, is defined by the gene expression programmes active in the cell. In this context, genetic elements with regulatory roles – i.e. affecting the expression of cohorts of target genes – might play an important function in setting the global biochemical and physiological properties of neurons, allowing for specific contributions to behaviour. We have recently identified several genes encoding small non-coding RNAs – i.e. microRNAs (miRNAs) – that are essential for a normal self-righting response in *Drosophila* (Picao-Osorio et al., 2015; Picao-Osorio et al., 2017; Issa et al., 2019). Several of these miRNAs – collectively termed SR-miRNAs (self-righting miRNAs; (Picao-Osorio et al., 2017)) – are expressed in the central nervous system, but others have been previously reported with expression in the peripheral nervous system (PNS) of larvae and adult (Pierce et al., 2008; Clark et al., 2010; Hilgers et al., 2010; Sun et al., 2012). Among these, we became particularly interested in *miR-263b* (Pierce et al., 2008; Hilgers et al., 2010) due to its pervasive evolutionary conservation (Pierce et al., 2008) (**Figure 2A-B**) and roles in multiple sensory and neural processes in the adult (Hilgers et al., 2010; Nian et al., 2019) and set to determine whether *miR-263b* might play a direct role in the SNs underlying self-righting behaviour. For this we first aimed at establishing the expression pattern of *miR-263b* in larval SNs. Fluorescent-activated cell sorting (FACS) experiments coupled to quantitative PCR expression assays (**Figure 2D-E**), demonstrate expression of *miR-263b* in Drosophila larval SNs. Furthermore, spatial expression analysis of a Gal4 insertion located within the *miR-263b* locus (Hilgers et al., 2010) confirms expression of this genetic element within SNs in the embryo (**Figure 2F**) providing independent experimental evidence that supports expression of *miR-263b* in SNs involved in self-righting control.

**Figure 2.**
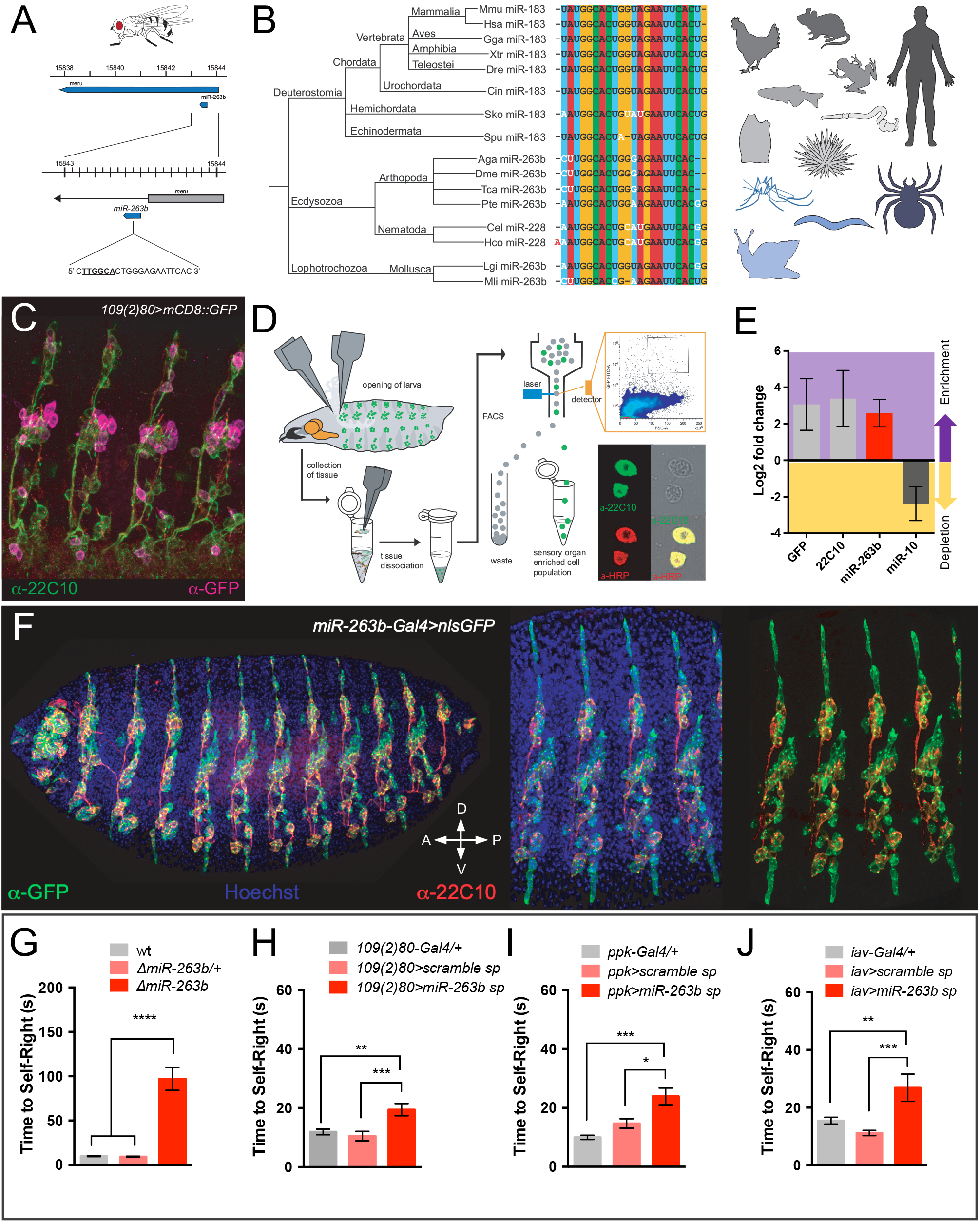
Expression and behavioural roles of *miR-263b*. **(A)** Genomic location of *miR-263b* in *Drosophila*. **(B)**. Evolutionary conservation of *miR-263b* throughout the animal kingdom. The sequence encoding *miR-263b* is highly conserved across large phylogenetic distances; there are several names for this miRNA: *miR-263b* in arthropods and molluscs; *miR-183* in deuterostomes; *miR-228* in nematodes. **(C)**. The driver line *109(2)80-Gal4* was used in cell sorting (FACS) experiments; its coupling to the *UAS-mCD8::GFP* construct labels larval sensory neurons (magenta). **(D)**. Schematic representation of the FACS protocol used to study miRNA expression in the sensory system. The process involved the dissection of larval tissue (top left), and cell type validation via antibody staining (bottom right). **(E)**. Relative gene expression of GFP and 22C10 sensory organ enriched cell population. **(F)** *miR-263b* expression using a the *miR-263b* driver confirms miRNA expression in sensory organ-associated cells in the *Drosophila* embryo. Sensory neurons are visualized in red (α-22C10) and expression of the *miR-263b* driver is depicted via GFP signal, in green (α-GFP). **(G)** *miR-263b* homozygote mutants (red) exhibit a pronounced delay in self-righting. **(H)** Reduction of *miR-263b* function via competitive inhibition (miRNA sponges) in multidendritic neurons and chordotonal neurons results in a very significant increase of self-righting time. **(I)** Functional inhibition of *miR-263b* in multidendritic (class-IV) neurons, or in **(J)** chordotonal organs, leads to significant delays in self-righting time. Altogether these results indicate that normal expression of *miR-263b* in the sensory system is necessary for normal larval behaviour.

The fact that *miR-263b* is expressed in larval SNs is consistent with a potential role of this miRNA in SNs; but falls short from demonstrating any functional roles. A more direct way to test the involvement of *miR-263b* in the biology of SNs is genetic removal: if *miR-263b* is essential for SN function, elimination of this genetic element from larvae is predicted to affect behaviours that rely on normal SNs, including self-righting. Experiments in **Figure 2G** show that homozygote as *miR-263b* mutant larvae display severe self-righting phenotypes demonstrating that this miRNA gene is essential for normal self-righting. Furthermore, functional attenuation of *miR-263b* by means of miRNA-specific sponges (Fulga et al., 2015) applied to the whole sensory domain or within specific sensory subsets (**Fig. 2H-K**) further confirms that *miR-263b*-mediated activities are necessary for normal self-righting behaviour to take place.

### miR-263b affects the physiology of sensory neurons

The regulatory nature of miRNAs makes them suitable for regulation of multiple functions within the organism, including developmental as well as physiological roles. To examine the point of action of *miR-263b* in the sensory system, we first set to establish whether this miRNA controls the development of SNs. For this we looked at the morphology of the sensory system in *miR-263b* null mutant larvae and wild type specimens. The complex and stereotyped morphology of the larval sensory field makes it particularly suitable as a system to examine effects on the developmental process. **Figure 3A-B** shows that detailed and systematic characterisation of the individual sub-components of the sensory system using confocal microscopy, in both, wild type and *miR-263b* null mutant larvae, reveals no detectable differences among these two genotypes (**Fig. 3B**).

**Figure 3.**
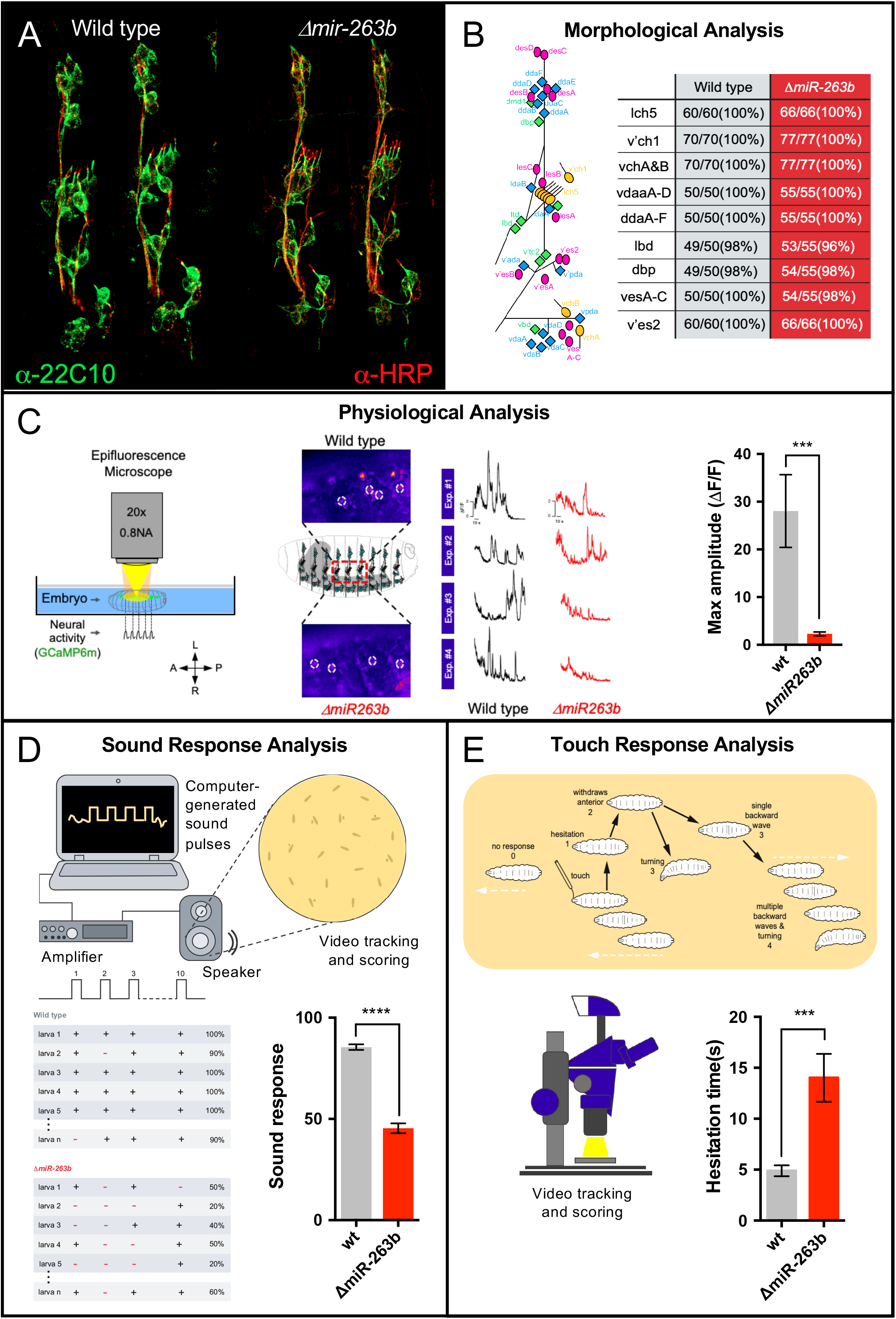
Effects of *miR-263b* on neural morphology, physiology and behaviour. **(A)** Labelling of sensory neurons (green, α-22C10) in stage 16 embryos in wildtype and *miR-263b* mutant specimens reveals the detailed morphology of the sensory system. **(B)** Comparison of the integrity and arrangements of a comprehensive series of sensory organ subtypes (see diagram on the left) in wildtype and *miR-263b* mutants; there are no morphological anomalies detected in *miR-263b* mutants. **(C)** Calcium sensor analysis (GCaMP6m) shows reduced neural activity in *miR-263b* mutants when compared to wildtype. **(D)** Experimental set up for sound response experiments (top). Diagram describing a representative set of sound response experiments in both genotypes (bottom left); quantification of responsiveness to sound (bottom right) is reduced in *miR-236b* mutants. **(E)** Experimental setup and scoring system for anterior touch-response experiments; video tracking and quantification of individual responses in normal and miRNA mutant larvae shows that *miR-236b* mutants exhibit an increased hesitation time (bottom right). These experiments suggest that *miR-263b* plays a role in the physiological control of sensory function with impact on behaviour.

The absence of morphological defects in the sensory system of *miR-263b* null mutants might indicate that this miRNA may impinge on the function (physiology) rather than the development of SNs. To test this possibility directly, we decided to quantify the spontaneous patterns of neural activity produced by SNs in miRNA mutants and compare these results to those obtained in normal specimens. For this we expressed a genetically-encoded calcium sensor (GCaMP6m) within the sensory system and visualised the resulting patterns of activity in live recordings. **Figure 3C** shows the results of these experiments where a significant decrease in neural activity patterns can be observed in *miR-263b* samples (**Fig. 3C**).

We reasoned that if *miR-263b* affects the general physiology of SNs, then other larval behaviours that rely on sensory input could also be affected in the absence of this miRNA. Two independent series of experiments confirm this. First, quantitative assessment of larval response to sound – previously shown to depend on sensory neurons, particularly, on chordotonal organs (Zhang et al., 2013) – reveal that *miR-263b* mutants display a marked decrease in sound responsiveness when compared to their wild type counterparts (**Fig. 3D-F**). Second, evaluation of the ability of larva to respond to touch (“anterior touch-response”) (Kernan et al., 1994) shows that *miR-263b* mutants display a reduced response with markedly pronounced “hesitation time” (**Fig. 3G-H**). Altogether, the combination of morphological, physiological and behavioural analyses of *miR-263b* mutants strongly indicates that this miRNA is required for normal sensory function in *Drosophila* larvae.

### A molecular model for miR-263b action in sensory neurons and self-righting behaviour

To advance the mechanistic understanding of the effects of *miR-263b* in the sensory system we decided to explore the potential points of action of this miRNA within SNs. Given that miRNAs are regulatory molecules (Bartel, 2018) their biological roles in the cell are likely to emerge indirectly, via effects on so-called “miRNA target” genes. Bioinformatic prediction of miRNA targets for *miR-263b* using the PITA and TargetScan algorithms (Kertesz et al., 2007; Agarwal et al., 2018) indicate a considerable number (n=186) of potential targets of this miRNA in the *Drosophila* transcriptome highlighted simultaneously by both algorithms; approximately one-third of these targets (28%) show expression in the nervous system (**Fig. 4A**). Within the predicted set of neural targets, 33% have demonstrated expression in the peripheral nervous system (PNS). Amongst the top-ten predicted *miR-263b* molecular targets with PNS expression, most genes encode factors with known functions in eye morphogenesis and differentiation, including: *WASp* (Wiskott-Aldrich syndrome protein) (Ben-Yaacov et al., 2001)), *tup* (tailup, also known as islet) (Thor and Thomas, 1997)), *Kr-h1* (Krüppel homolog 1) (Fichelson et al., 2012)), *arr* (arrow) (Wehrli et al., 2000)), *sca* (scabrous) (Mlodzik et al., 1990), *repo* (reverse polarity) (Xiong et al., 1994)), *Vav* (Vav guanine nucleotide exchange factor) (Malartre et al., 2010)) and the basic helix-loop-helix (bHLH) transcription factor Atonal. The latter was of particular interest to us due to its role as a proneural gene during the specification and development of chordotonal organs (Jarman et al., 1993);(Jarman and Ahmed, 1998), amongst other sensory organs (Jarman et al., 1995; Gupta and Rodrigues, 1997). Chordotonal organs are internal stretch receptors that detect cuticle movement and vibration, and have been shown to be important for detecting sound (Zhang et al., 2013). As we expect chordotonal organs to play roles in all three behaviours tested (i.e. self-righting, sound-and touch-response), we selected *atonal* as an entry point for detailed functional studies.

**Figure 4.**
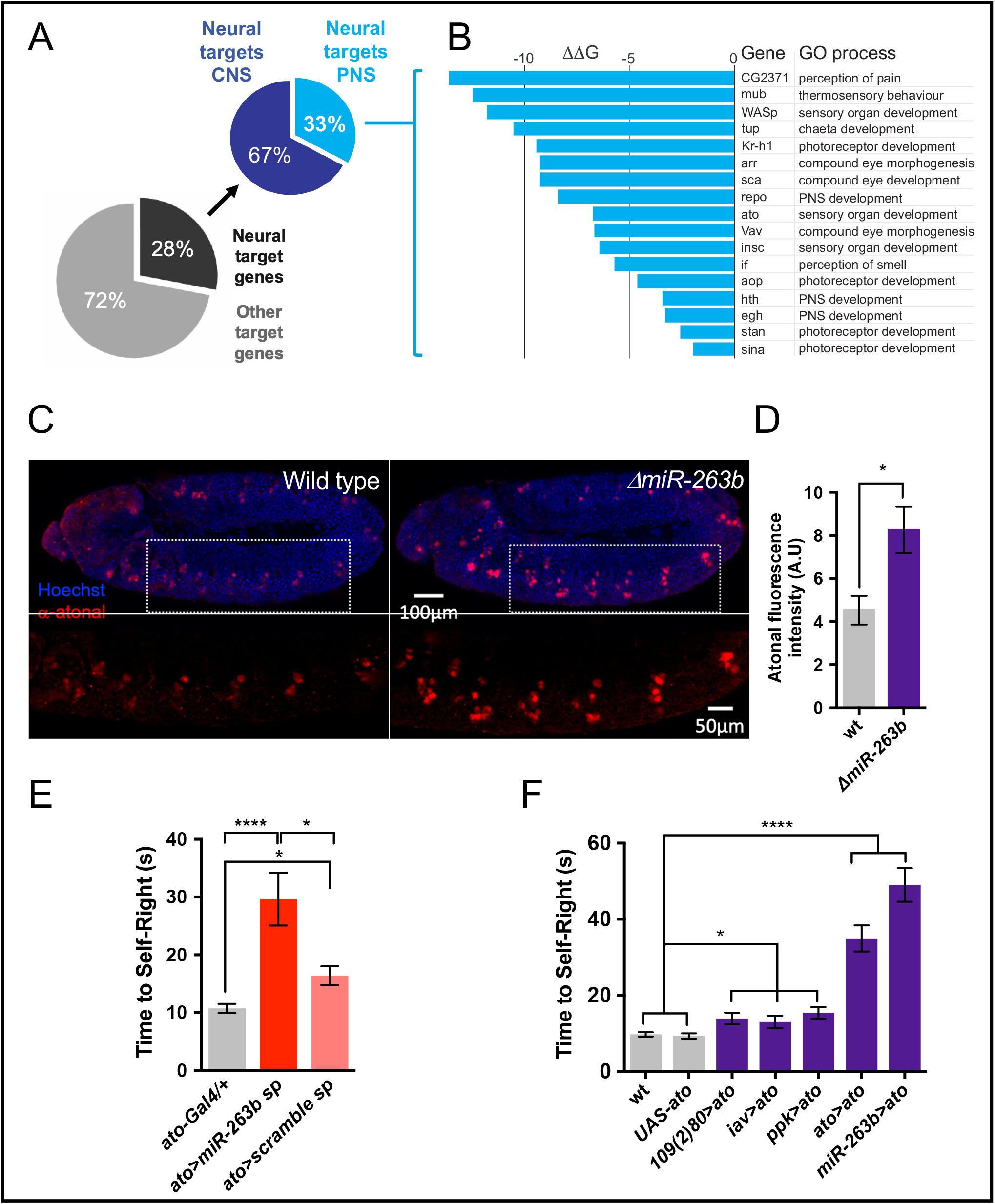
Potential molecular targets of *miR-263b* and their roles in self-righting. **(A)** Pie charts representing shared predicted target genes of *miR-263b* through the combined output of the PITA and TargetScan algorithms. Out of 186 total miRNA targets highlighted by both computational approaches, 52 (28%) can be classified (GO terms) as “neuronal targets”; within this group 17 relate to the peripheral nervous system (PNS). **(B)** Details on the 17 predicted target genes involved in the PNS, note the pro-neural gene *atonal* (*ato*) within the top-ten predicted target genes of *miR-263b*; **(C)** Atonal protein expression analysis in wildtype and *miR-263b* mutant specimens reveals that Atonal expression is elevated in *miR-263b* mutants; **(D)** Quantification of antibody labelling experiments for Atonal demonstrate that the levels of this proneural protein are significantly increased in *miR-263b* mutants **(E)** Reduction of *miR-263b* function within the *atonal* expression domain increases self-righting time. **(F)** Artificial increase of *atonal* expression (i) within different sensory fields (activated via *109(2)80*, *ppk* and *iav* drivers), (ii) within the endogenous *atonal* expression domain, or (iii) within the *miR-263b* transcriptional domain all result in significantly increased self-righting times. These data suggest that atonal might be one of the molecular targets that mediates the roles of *miR-263b* in the sensory system.

Three independent series of experiments support the model that *atonal* is one of the factors that mediates the actions of *miR-263b* in the sensory system with effects on self-righting control. First, *atonal* is expressed in sensory neurons, and its expression is increased in miRNA null mutants, consistently with a de-repression effect caused by removal of a repressive regulatory miRNA (**Fig. 4C-D**). Second, expression of a *miR-263b-sponge* within the *atonal* domain is sufficient to trigger a self-righting phenotype (**Fig. 4E**). Third, artificial upregulation of *atonal* within its natural transcriptional domain in wild type larvae – so that its expression level emulates those observed in *miR-263b* mutants – is sufficient to phenocopy the self-righting defect observed in miRNA mutants (**Fig. 4F**); similar effects are observed when atonal is specifically delivered within sub-field of the sensory system or when driven within the transcriptional domain of *miR-263b* (**Fig. 4F**). Altogether, these results suggest a model by which *miR-263b*-mediated repression of *atonal* in sensory neurons is essential for a normal self-righting response.

## DISCUSSION

We have employed *Drosophila* larvae to investigate the sensory elements required for triggering an evolutionarily-conserved 3-D postural behaviour (self-righting) and established that normal expression of a miRNA gene, *miR-263b*, in the larval sensory system is necessary for a normal self-righting response. Our experiments also explore the causal links between the absence of *miR-263b* and the resulting behavioural effects, and show that absence of *miR-263b* does not seem to affect the complex morphological organisation of sensory neurons in the fruit fly larvae. Instead, we observe an impairment in several sensory functions (touch-response, sound detection) as well as a reduced level of spontaneous neural activity in the sensory neurons of *miR-263b* mutants. Based on these results we suggest that *miR-263b* is required for the normal physiological control of sensory function in *Drosophila* larvae, with no detectable effects in the formation of the sensory system. At the mechanistic level, the combination of bioinformatic, gene expression and behavioral analyses indicate that *miR-263b* might exert its actions – at least in part – through repression of the bHLH transcription factor *atonal*, a regulatory proneural gene with known roles in the formation of sensory elements in *Drosophila* (Jarman et al., 1993; Jarman et al., 1995; Bossuyt et al., 2009) and mammals (Bermingham et al., 1999; Zheng and Gao, 2000)

Evidence from behavioural genetic studies across different species indicates that the genetics of behaviour is often complex, and that the path between individual genes and behavioural traits is frequently intricate and hard to work out (Greenspan R.J., 2008). Here we see a different picture, in which a reduction in the expression of a miRNA gene (caused by either a single loss-of-function mutation or tissue-specific functional attenuation) leads to a consistent behavioural change. Yet, how exactly a reduced level of expression of *miR-263b* modifies the properties of sensory elements in the fruit fly maggot thus causing a change in behaviour, remains open. Our observations, nonetheless, suggest that *miR-263b* absence leads to an increase in the expression of *atonal*, which, instead of derailing the developmental process, affects the biochemistry and physiology of sensory cells. Indeed, experiments in rat explants indicate that an excess of *atonal*’s orthologue in the rat (Math1) can promote the formation of hair cells out of a population of postnatal utricular supporting cells (Zheng and Gao, 2000); this shows that going above a particular ‘set-point’ of atonal protein concentration (as observed in our study) is sufficient to modify the cellular processes in sensory elements. Given that *atonal* is expressed in *Drosophila* chordotonal organs (Jarman et al., 1993; Jarman and Ahmed, 1998) as well as in the cochlear system of mammals (Bermingham et al., 1999; Yang et al., 2012; Jarman and Groves, 2013), it might be plausible that *atonal*-dependent events could impact on behaviour in other systems too (see below).

Interestingly, a previous investigation in our laboratory (Picao-Osorio et al., 2015) showed that a deficit of another miRNA, *miR-iab4*, caused a change in the physiology of larval motor neurons involved in self-righting. In this case, *miR-iab4* mediates the repression of the Hox protein Ultrabithorax (Ubx) – a homeodomain-containing transcription factor (Bridges C.B., 1923; Sánchez-Herrero et al., 1985; Mallo and Alonso, 2013); in turn, the increase of *Ubx* phenocopies the behavioural effects observed in *miR-iab4* mutants (Picao-Osorio et al., 2015). Building on these examples, we would like to suggest the model that miRNA-dependent control of behaviour relies on maintaining the expression levels of a small set of transcription factors within a particular concentration range; departures from such range may lead to gene regulatory effects with impact on neuronal biochemistry, in ultimate instance, modifying the physiological properties of the cell. However, current views on the molecular mechanisms of miRNA function strongly indicate that miRNAs can regulate multiple targets within the cell (Bartel, 2018; McGeary et al., 2019) arguing that regulation via single or few target genes is unlikely. A potential framework that reconciles both scenarios is that for specific cells (i.e. neurons) miRNA-dependent regulation of just a small subset of targets (e.g. *ato, Ubx*) is critical for their biological function, while other regulatory events are simply tolerated or compensated by the regulatory networks of the cell (Alonso, 2012).

Our results show that ablation of either all, or genetically-defined subtypes of sensory elements in the larvae lead to a significant impairment in self-righting, but how sensory information is transformed into the actual slef-righting response in the larva is still unknown. A current effort (O’Garro-Priddie, Picao-Osorio, Cardona, and Alonso, *in preparation*) is using neural connectomics and reconstruction at synaptic resolution to map the neural substrates of self-righting behaviour and should – when completed – offer a cellular platform for the investigation of how information in the sensory system is transmitted and converted into the motor patterns that underlie the self-righting sequence.

We note that *miR-263b* is evolutionary conserved across large taxonomical distances. Building on this, and the similarly broad conservation of *atonal* (Quan and Hassan, 2005; Cai and Groves, 2015) and that of self-righting itself (Ashe, 1970; Faisal and Matheson, 2001; Jusufi et al., 2011), we speculate that *miR-263b* and *atonal* orthologues in other species may play a role in self-righting and other postural control mechanisms in other orders of animals. Although the idea that the same set of miRNAs and their targets may control complex behavioural responses in animals with drastically distinct body plans such as flies and mammals, seems highly unlikely, or even absurd, a recent observation in our lab may help keeping this plausibility open: *miR-iab4*-dependent control of *Ubx* expression is required for normal self-righting behaviour in both *Drosophila* larvae and adults, two morphs of the fly that bear radically different morphologies, neural constitution and biomechanics.

Our study shows that the function of sensory neurons may rely on the normal expression of small non-coding RNAs, and that this is relevant for a complex and adaptive behavioural sequence in *Drosophila* larvae illustrating that behaviour emerges from a careful balance in the expression of transcriptional as well as post-transcriptional gene regulators within the nervous system.

## ACKNOWLEDGEMENTS

We thank members of the Alonso lab for helpful discussions and comments. We wish to thank Sofia Pinho, Tom Baden, and Virginia Mahou at *Sussex Neuroscience* for their assistance with dissections, functional imaging and behavioural experiments. We also thank Jonathan Wing at the *Genome Damage and Stability Centre* at Sussex for his help with FACS experiments. We also thank Sherry Aw and Daniel Marenda for sharing fly strains and antibodies, respectively. This research was funded by a Wellcome Trust Investigator Award made to C.R.A. (098410/Z/12/Z).

## MATERIALS AND METHODS

### *Drosophila melanogaster* strains

Cultures of *D. melanogaster* stocks were kept under standard conditions with 50-60% relative humidity and a 12h light/dark cycle at 18 ◻C, while working copies were held at 25 °C. The following stocks were used in this study: 109(2)80-Gal4 (BDSC #8769), iav-Gal4 (BDSC #52273), ppk-1.9-Gal4 (stock described in Ainsley et al. 2003 (Ainsley et al., 2003), gift from Matthias Landgraf), ato-Gal4 (BDSC # 6480), 109(2)80-Gal4,UAS-mCD8::GFP (BDSC #8768) ΔmiR-263b-Gal4 (stock described in Hilgers et al. 2010 (Hilgers et al., 2010), gift from Sherry Aw), UAS-TeTN Lc (BDSC #28838), UAS-nls-GFP (BDSC #4775), UAS-miR-263b sponge (BDSC #61403), *UAS-scramble sponge* (BDSC #61501), UAS-GCaMP6m (BDSC #42748), *ΔmiR-263b* (BDSC #58903), UAS-ato (BDSC #39679); *w^1118^* flies (BDSC #5905) were used as controls.

### Immunohistochemistry

Embryo collection, dechorionation, devitellinization and fixation followed the standard protocols described and samples were stored in 100% Methanol at −20 °C. To obtain various embryonic stages an over-night collection was used. The samples were gradually rehydrated, rinsed in PBTx (1x PBS, 0.3% Triton X-100) and washed with PBTx 4x 30min. Subsequently, the samples were incubated with primary antibody (in PBTx) over night at 4 °C. Primary antibodies and their concentration used are: 1:10 mouse anti-22C10 (Developmental Studies Hybridoma Bank), 1:2000 chicken anti-GFP (Abacam Probes), 1:100 guinea pig anti-atonal (gift from Daniel Marenda) and goat anti-HRP-A556. Primary antibody was removed with 3 rinses and 4x 30 min PBTx washes. The secondary antibody was added and incubated for 2h at room temperature. Secondary antibodies and their concentrations used are: 1:500 anti-mouse Alexa Fluor 488 (Invitrogen), 1:500 anti-mouse Alexa Fluor A555 (Invitrogen), 1:500 anti-chicken Alexa Fluor 488 (Invitrogen), 1:500 anti-guinea pig Alexa Fluor A555 or A488. During the incubation with the secondary antibody 1:500-1:1000 Hoechst 33342 (Life Technologies) was added. Samples were rinsed 3 times, washed 4x 30 min with PBTx and transferred to 75% glycerol for mounting. Samples were imaged on a Leica SP8 confocal laser scanning microscope and further processed using Image J/FIJI and Adobe Photoshop CS6. Schematic representations and figure arrangement were made with Adobe Illustrator CS6.

### *In Vivo* Calcium Imaging

The calcium sensor GCaMP6m was used to measure the Ca2+ signal in SNs. Calcium imaging were conducted on stage 17 embryos to reduce movement during the recording. Parental lines were raised at 25°C in collection cages bearing apple juice-based medium agar plates, supplemented with yeast paste. Prior to recording, stage 16 embryos were collected, dechorionated and transferred into a drop (1μL) of PBS (to prevent dehydration) previously left on a Poly-L-Lysine coated glass slides. The embryo was placed with its lateral side up, to allow visibility of the majority of SNs. Next, the Ca2+ signal within the SNs was captured during 3-min using Leica DM6000 microscope (Leica Microsystems) and processed with the software *Fiji image J*. GCaMP signals from the soma were analysed. The average signal from the first 10-s was taken as fluorescence baseline F_0_ to calculate the ΔF/F_0_ for each recording.

### Cell sorting experiments

The FACS dissociation protocol followed was the one described by Harzer and collegues (Harzer et al., 2013) with some adjustments. Control (w^1118^) and experimental larvae (109(2)80-Gal4,UAS-mCD8::GFP; 50h old) were opened anterodorsally and the gut was partly removed. For each genotype larvae were dissected for 30 min, usually obtaining 50-60 larvae, which were pooled in an Eppendorf tube filled with Rinaldini’s solution (800mg NaCl, 20 mg KCl, 5 mg NaH_2_PO_4_, 100 mg NaHCO_3_, 100 mg Glucose in 100 ml H_2_O). The larvae were washed once with 500 μL Rinaldini’s solution. Rinaldini’s solution was replaced with dissociation buffer, the mixture was incubated for 1h at 30°C and gently mixed twice during incubation. Dissociation buffer needs to be prepared fresh, by adding 25 μL Collagenase Type I (20 mg/mL, Sigma-Aldrich) and 25 μL Papain (20 mg/mL, Sigma-Aldrich) to 200 μL complete Schneider’s culture medium (5 mL heat inactivated fetal bovine serum, 0.1 mL Insulin, 1 mL PenStrep, 5 mL L-Glutamine, 0.4 mL L-Glutathione and 37.85 mL Schneider’s medium). All subsequent washing steps need to be carried out very slowly and carefully in order to avoid premature dissociation of the larval bodies. After the removal of the dissociation solution, the samples were washed twice with 500 μL Rinaldini’s solution first, and then twice with 500 μL complete Schneider’s culture medium. All medium was removed and the larval bodies were dissociated using 200 μL complete Schneider’s medium, which was pipetted up and down (medium and larvae) with as little foaming as possible. The solution started to appear homogenous after pipetting was repeated about 50 times. The cell suspension was filtered through a 30 μm mesh into a 5 mL FACS tube, placed on ice and immediately taken to a FACS system (BD-FACSMelody™). One biological replicate comprised around 3,000 sorted cells (one sorting event); cells were sorted directly into TRIzol (Invitrogen) and stored at −80°C until further processing.

### Quantitative PCR assays

A standard phenol/chloroform protocol was employed to extract RNA from FACS samples using Phase Lock Gel 2mL Heavy tubes (Fisher Sceintific). RNA was treated for 30 min with DNAse (TURBO DNA-free ™, Invitrogen) subsequently. SuperScript® III First Strand (Invitrogen) was used for cDNA synthesis. A single qPCR reaction is made of 5 μL 2xSYBR green mix (LightCycler® 480 SYBR Green I master, Roche), 2 μL H_2_O, 1 μL 5 μM forward primer, 1 μL 5 μM reverse primer, 1 μL cDNA (diluted 1:2 with H_2_O). Every sample was run in technical triplicates. The cycle conditions used were 40x (10 sec 95 °C, 20 sec 60 °C, 20 sec 72 °C) with fluorescent readings during annealing and elongation on a QuantStudio 3 machine (Thermo Fisher). Three reference genes were tested, Cdc5, RpS9, and eEF1◻, and Cdc5 was chosen as appropriate reference gene. Primer sequences used are listed below.

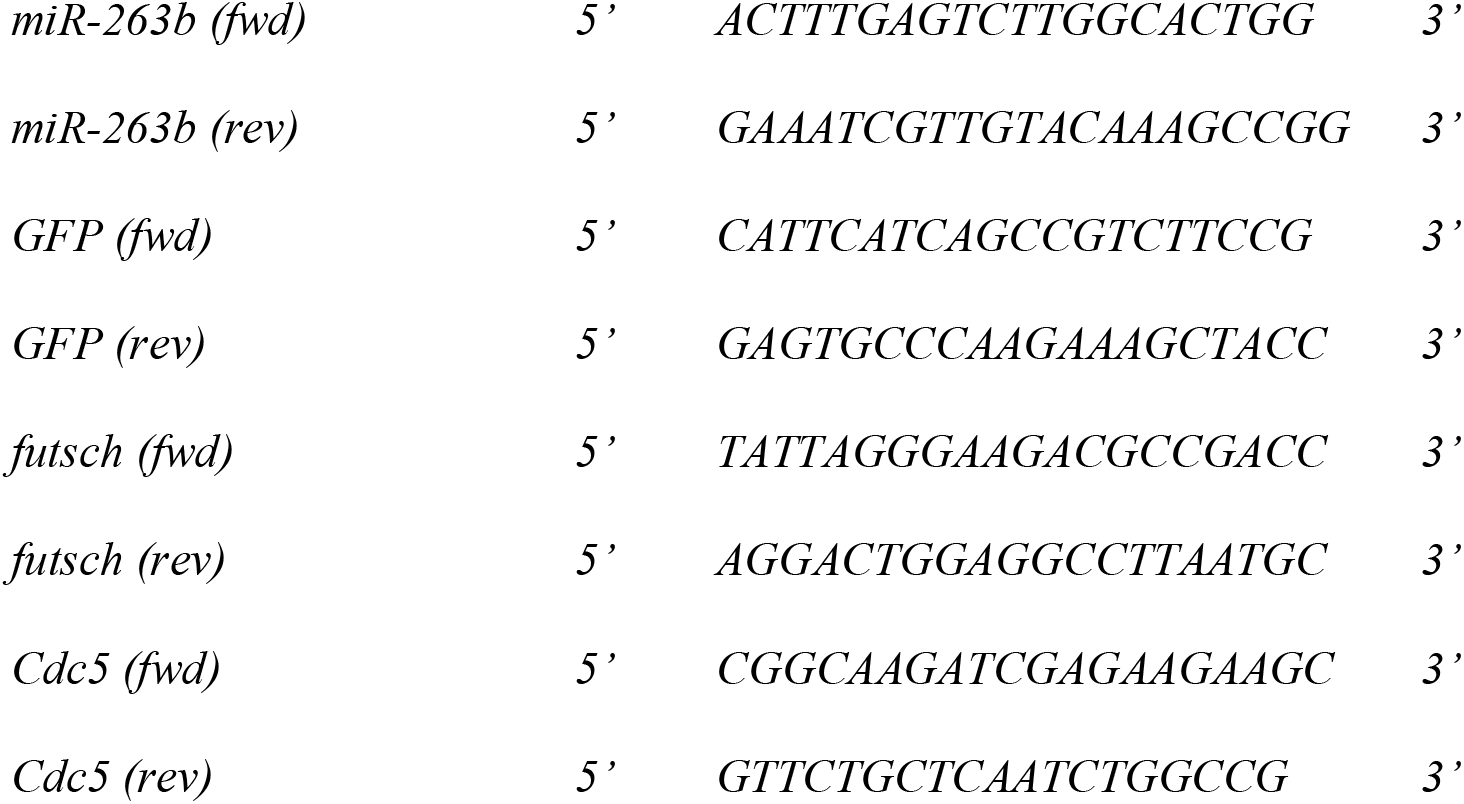

For each primer set a standard curve was generated using serial dilution of the cDNA (undiluted, 1:2, 1:4, 1:8, 1:16 and 1:32), which was employed to calculate primer efficiency E (E=10^(−1/slope)^). The primer efficiency was incorporated into the formula to calculate fold change values (R) as described in (Pfaffl 2001):

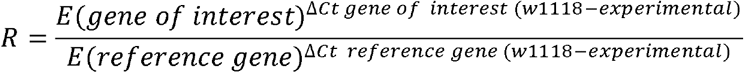

### Sequences, statistics and miRNA target prediction

Sequences for miR-263b/183/228 were recovered from miRBase.org (release 22) for schematic representation of the phylogenetic tree. Statistical analysis was done using Microsoft Excel. Statistical significance was calculated using Student’s t-tests with a *p* value threshold of 0.05 for significance. For *atonal* expression, fluorescent signal comparisons from wildtype and *ΔmiR-263b* mutant embryos were processed the same/at the same time and images were collected using the same settings. Image J was employed to analyse fluorescent intensity of individual *atonal*-positive clusters, with the help of the ROI manager. A ROI was manually drawn around an *atonal*-positive cell cluster as well as a region adjacent to the *atonal*-positive clusters where *atonal* is not expressed (to normalize background expression). This was repeated for a minimum of 7 clusters (14 cluster-pairs), all located in the thoracic or abdominal region of the embryo. After all ROIs were selected, they were measured using “Measure” button within the ROI manager tool. Per ROI 4 values related to fluorescence were provided by the program: area, mean, minimum and maximum. First, mean intensity was calculated per area (intensity/area). To calculate the fluorescent intensity of an *atonal*-positive cluster, the intensity/area of the background expression (*atonal*-negative area) was subtracted from the intensity/area of the *atonal*-positive cluster. Per embryo between 7-10 *atonal*-positive clusters were measured and the average was taken. At least 10 embryos were analysed per genotype. For *D. melanogaster miR-263b* target prediction we used PITA (Kertesz et al., 2007) and TargetScanFly 7.2 (Agarwal et al., 2018).

### Behavioural tests

All embryos/larvae were kept on apple juice agar plates containing a small amount of yeast paste at 25 °C. Self-righting and anterior touch response assays were carried out on freshly hatched L1 larvae (less than 30 min old). For those, late stage embryos were transferred to a fresh apple juice agar plate (without yeast paste) and monitored for emerging larvae. Newly hatched L1 larvae were transferred to another fresh apple juice agar plate, which was used for testing throughout one session (1 biological repeat). Early L3 larvae (72h) were used for the sound response assay (startle assay). At least 3 biological replicates with a minimum of 20 larvae per replicate were analysed for each behavioural test. A paint brush was used to roll larvae for self-righting tests, which were otherwise performed as described in (Picao-Osorio et al. 2015). An eyelash was employed to deliver a soft stroke to the anterior region (head and thorax only) for anterior touch response assays. The protocol followed those described by Kernan and colleagues (Kernan et al., 1994) with slightly altered scoring, for scheme see Fig. 3G. Overall response score and hesitation time were analysed. For sound response/startle assays the protocol developed by Zhang and colleagues (Zhang et al., 2013) was followed. In brief, 72h larvae were washed with PBS and transferred to a testing plate. The plate was put on top of a speaker and videotaped from above. Larval response to sound was assayed by stimulation with a 1 sec sound pulse (pure tone, 400 Hz), which was repeated 10 times.

